# Rhythm of The Night (and Day): Predictive metabolic modeling of circadian growth in *Chlamydomonas*

**DOI:** 10.1101/2022.01.24.477634

**Authors:** Alex J. Metcalf, Nanette R. Boyle

**Affiliations:** Chemical and Biological Engineering, Colorado School of Mines, 1613 Illinois St. Golden CO 80401

**Keywords:** transcriptomics, systems biology, algae, diurnal light, metabolic modeling

## Abstract

Algal cells experience strong circadian rhythms under diurnal light, with regular changes in both biomass composition and transcriptomic environment. However, most metabolic models – critical tools for bioengineering organisms – assume a steady state. The conflict between these assumptions and the reality of the cellular environment make such models inappropriate for algal cells, creating a significant obstacle in engineering cells that are viable under natural light. By transforming a set of discreet transcriptomic measurements from synchronized *Chlamydomonas* cells grown in a 12/12 diel light regime (1) into continuous curves, we produced a complete representation of the cell’s transcriptome that can be interrogated at any arbitrary timepoint. We clustered these curves, in order to find genes that were expressed in similar patterns, and then also used it to build a metabolic model that can accumulate and catabolize different biomass components over the course of a day. This model predicts qualitative phenotypical outcomes for the *sta6* mutant, including excess lipid accumulation (2) and a failure to thrive when grown diurnally in minimal media (3), representing a qualitative prediction of phenotype from genotype even under dynamic conditions. We also extended this approach to simulate all single-knockout mutants with genes represented in the model and identified potential targets for rational engineering efforts.

**SIGNIFICANCE STATEMENT:** We have developed the first transient metabolic model for diurnal growth of algae based on experimental data and capable of predicting phenotype from genotype. This model enables us to evaluate the impact of genetic and environmental changes on the growth, biomass composition and intracellular fluxes of the model green alga, *Chlamydomonas reinhardtii*. The availability of this model will enable faster and more efficient design of cells for production of fuels, chemicals and pharmaceuticals.

## INTRODUCTION

The dawn of a new century has also led to an awakening about our energy use; there has been a concerted push to develop more sustainable sources of energy and feedstock chemicals. One of the major challenges in developing more sustainable fuels and chemical supply chains is that the current industry is large and expensive capital has already been spent to develop the infrastructure for processing and distribution. To avoid an extensive and expensive redesign of infrastructure, renewables will have to be chemically similar to the fossil fuels we currently rely on. The fossil fuels we use today are derived from prehistoric biomass; therefore, it is a logical extension to engineer biomass to produce high concentrations and/or yields of equivalent chemical feedstocks and fuels that can be ‘dropped in’ to current infrastructure. This approach has been successfully commercialized for a handful of chemicals, such as 1,3 propanediol (4, 5), 1,4-butanediol (6–9), and succinate (10) (amongst others). One drawback of these commercialized ventures is that they utilize heterotrophic bacteria, which require a source of reduced carbon which is derived from corn or sugar cane. A more sustainable approach is the use of photosynthetic organisms that can grow on CO_2_; however, the development of photosynthetically derived fuels and chemicals has fallen far short of the touted potential because the unique demands of photosynthetic organisms have not been considered in the strain development process. For example, algae and cyanobacteria have extremely strong circadian rhythms (11–13) which control the expression of cellular transcripts, proteins, and metabolic fluxes (1, 12, 14). Additionally, they are strongly entrained – cells that are removed from light/dark cycles will continue to display circadian behaviors for days afterward (15). These rhythms have a profound impact on the cell but unfortunately, they have not been studied well in the context of how they impact engineering efforts. Almost all metabolic engineering efforts in algae and cyanobacteria rely on growth in laboratory conditions with a continuous supply of light (16–29), but it is economically prohibitive to provide constant artificial light. If we are to grow these organisms outside in the most cost-effective manner, natural circadian rhythms must be considered when designing production strains.

Metabolic modeling techniques (especially constraints-based stoichiometric models) have proven extremely useful in minimizing the time it takes to develop heterotrophic production strains (30–33). One of the main assumptions in stoichiometric modeling approaches is steady state; this means that metabolite pool sizes are assumed to stay constant. While this approach works well for heterotrophic bacteria in exponential growth, it cannot be used to model the transient nature of photosynthetic growth in day/night cycles. When grown in diurnal light, 30 – 50% of genes show cyclic expression in *A. thaliana* (34, 35) and up to 80% of genes in cyanobacteria, diatoms and alga show periodic expression (36–39). These changes in gene expression impact metabolic fluxes, so to build a predictive model of photosynthetic metabolism, changing gene expression must be integrated into the model. Circadian rhythms don’t only affect the cell’s fluxes, the actual biomass composition of the cell changes over the course of the day as well; photosynthetically grown cells store starch or glycogen during the day to support metabolism at night. This also requires a different formulation of stoichiometric modeling as the steady state assumption necessitates the use of a static biomass formation equation, but diurnally grown cells alter their biomass composition significantly from day to night. The unique properties of circadian growth in photosynthetic cells means that a new modeling framework must be developed in order to be able to more accurately predict metabolism.

Here, we describe the development of a transient modeling approach capable of predicting phenotype from genotype in the model green alga *Chlamydomonas reinhardtii*. This model combines the easy to implement stoichiometric constraints-based model of *C. reinhardtii* (40) with experimentally determined transient genetic constraints on reaction bounds and a decoupled biomass formation equation that enables flexibility in the production of biomass components. By incorporating the additional complexity to account for the transient nature of diurnal growth, we are able to increase the predictive nature of our model and simulate intracellular fluxes, biomass composition and growth in diurnal light.

## RESULTS AND DISCUSSION

### Fitting and Clustering of Transcriptomic Profiles

Prior to using the experimental data for constraining the model, it first has to be converted from discrete data to continuous so that we can model smaller time steps than the experimental data (2-hour time steps). This way, the transcript abundance of any transcript can be modeled at any time step. By fitting transcripts to best fit functions, we can also easily compare the profile of expression across several genes (Figure 1). The transcript expression profiles are clustered with other transcripts of similar expression patterns. We have chosen to highlight 4 specific genes and the most closely associated expression profiles in Figure 1. As previously discussed, up to 80% of genes in cyanobacteria, diatoms and alga show periodic expression (36–39) in diurnal light, which makes it difficult to identify housekeeping genes to normalize gene expression to. Therefore, the first gene we investigated further was *RACK1* (*CBLP*), which is commonly used as a housekeeping gene (41), which shows very little variation in expression over time. We also found 19 named genes that are clustered closely with *RACK1*; these may be good candidates for housekeeping genes (see supplemental file 2). We noticed that the expression profile of many genes was most accurately modeled by the Kronecker Delta function, when we investigated this cluster, we saw that they cluster with known stress response genes, such as *LHCSR3.1* (42) (see supplemental file 3). This is likely more an artifact of the experimental design, where the light is turned on immediately at the onset of day instead of ramping up as it would do in natural conditions. It is well known that Ribulose-1,5-bisphosphatase carboxylase-oxygenase (RuBisCO) is a light regulated enzyme (43); our expression profile of *RBCS1*, the gene encoding the small subunit of RuBisCO, the shows that the cell anticipates the onset of light by starting transcription of this gene prior to the onset of light and the peak transcript abundance occurs 6 hours after the onset of light. This profile is also clustered with multiple other named genes, including *FUO1*, associated with the electron transport chain, and *PGK1*, which encodes for phosphoglycerate kinase, a critical enzyme within the Calvin cycle. (see supplemental file 4 for more details). We also looked into genes known to be expressed at night, such as *FAP85*, a gene encoding a flagellar protein; genes associated with the synthesis of cilia are coordinately expressed at night following cell division (44). Other genes closely clustered with *FAP85* include *CPC1*, which is associated with the central pair microtubule complex (45), as well as ten other genes specifically identified as encoding flagella associated proteins (see Supplemental File 5). Fitting transcript profiles to continuous functions not only aides in the development of transient constraints for the model, but it also helps to identify similar expression profiles and potentially gene function.

**Figure 1.**
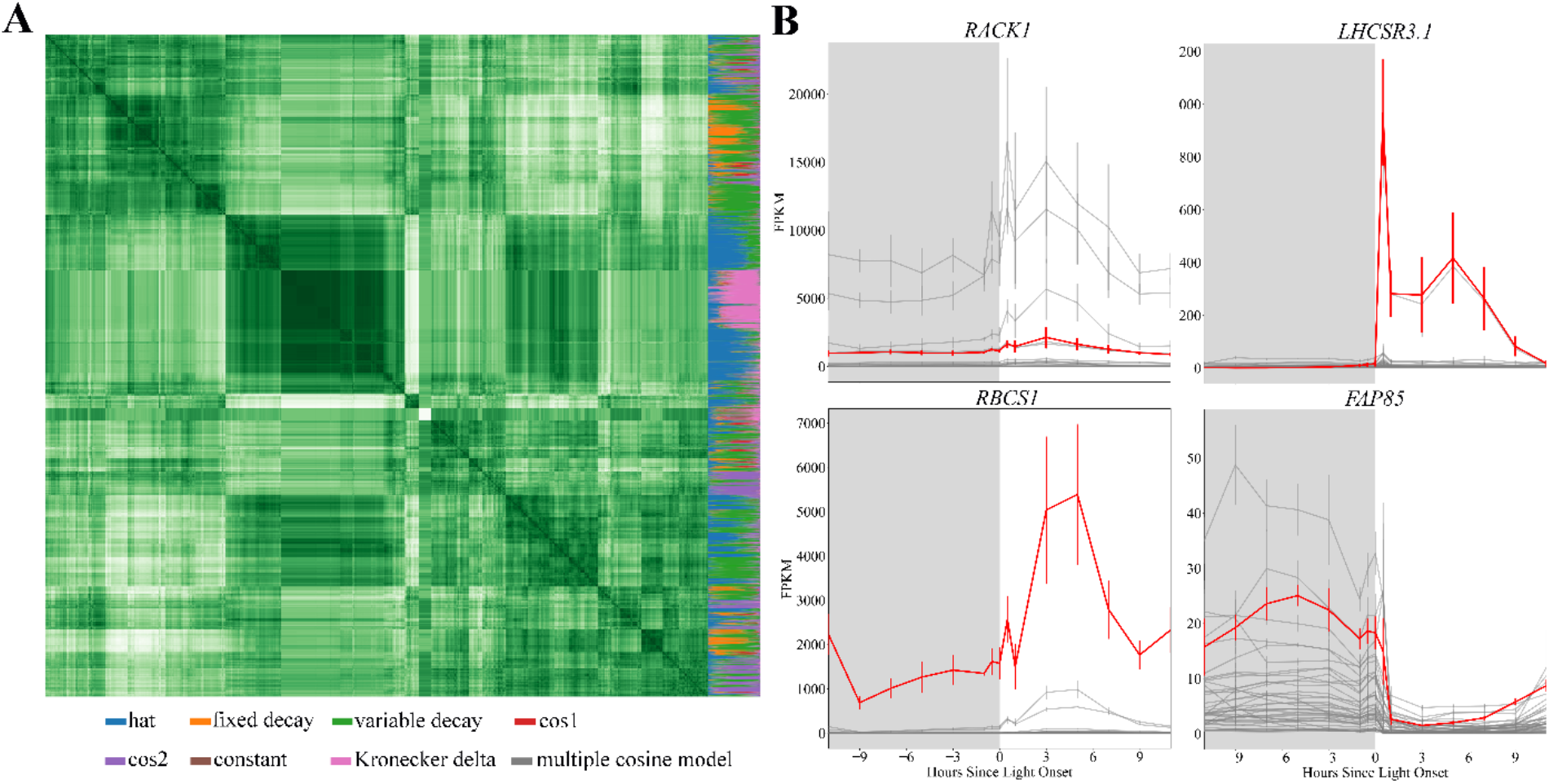
Clustering time dependent transcriptomic data provides insight into gene expression patterns. A) Heat map of all expression patterns within the dataset. Every pixel in the main square of this image represents the degree of closeness between a pair of best-fit transcript functions, while the column on the right side indicates the relative weight of each of the best-fit models to the transcript in question; the larger amount of a given color in the column, the higher probability that that model fits the actual expression pattern better than any other of the evaluated choices. The multicolored column on the right of the heat map indicates the weight of each proposed curve fit to the gene expression pattern for a given gene; for example, if the majority of the column is pink, then that gene is best fit by the Kronecker Delta function. B) Examples of select genes of interest (shown in red) and genes which are most closely co-expressed (dark grey lines). The genes included in each cluster shown above are provided in supplemental files 2-5.

### Dynamic Diel Modeling

By dynamically adjusting the model constraints according to transcriptomic abundance, we constructed a metabolic model that is capable of accurately describing the shifts in metabolism due to circadian rhythms and changing light conditions. In our simulations, we had an on/off onset of light but due to the addition of gene expression constraints we see a ramp up of carbon flux though the Calvin Benson Bassham Cycle over the course of the day; it peaks at 5 hours, after which carbon flux is ramped back down as the simulation moves toward the onset of night (Figure 2, animated in Supplemental File 6). Despite the simulated cell experiencing the same amount of light at both timepoints, it utilizes less of it early in the day because the transcripts associated with the light reactions are less present, thereby reflecting transient physical limitations within the constraints of a dynamic model. These daytime ramps are driven by changes in expression of the photosystems and associated photosynthetic light reactions, but they end up reflected in the carbon flux throughout the cell as well (see Supplemental Files 5-18). The cell shuttles large amounts of carbon between the chloroplast and the mitochondria, likely to make use of the more efficient mitochondria ATP synthase (46). The cell also experiences a night slowdown, during which time it simply attempts to produce the maintenance ATP and does not encounter any significant reaction bounds. This phenomena is supported by the finding of Strenkert et al. that the cell does not utilize its full respiratory capacity at night (1).

**Figure 2.**
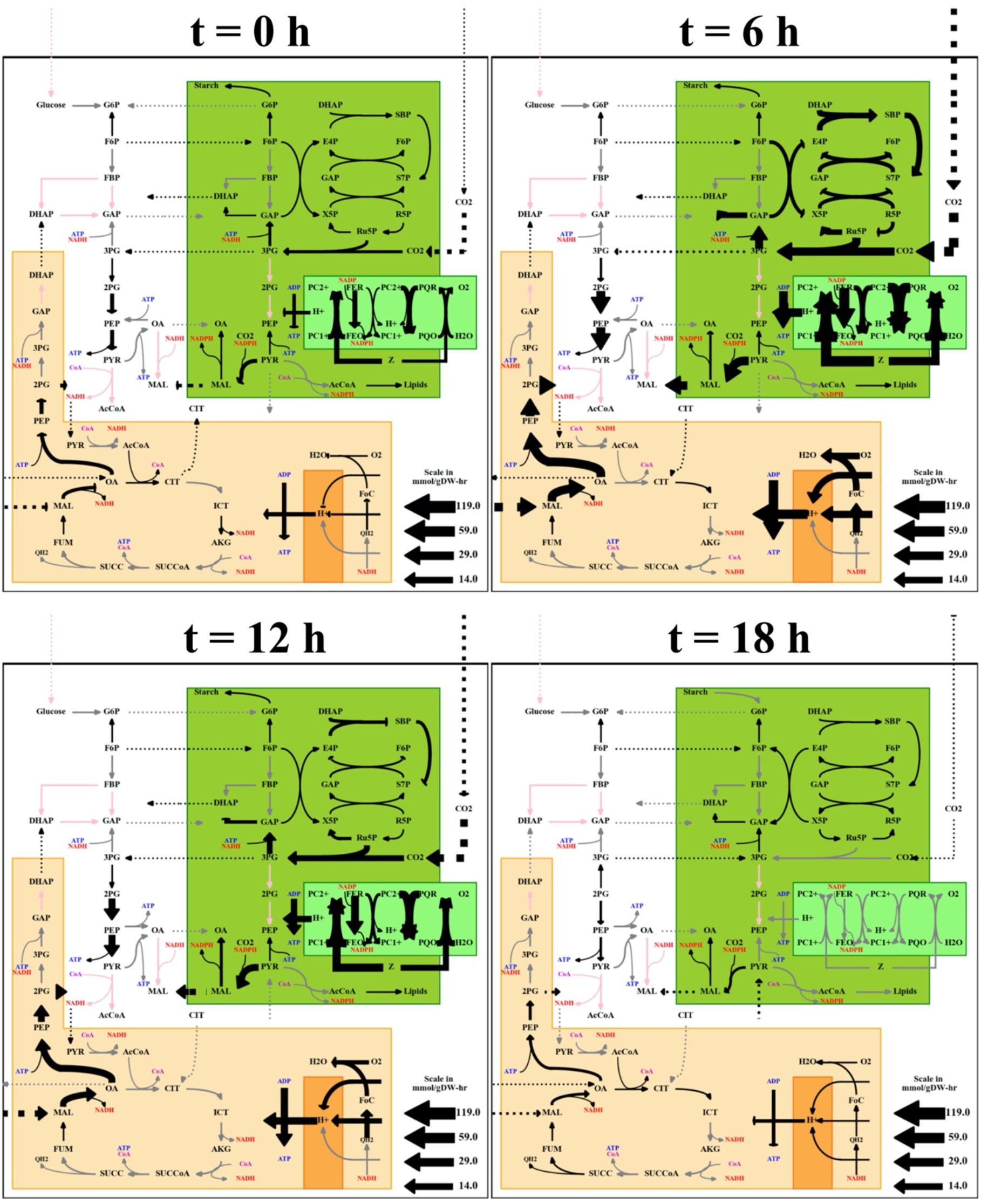
Flux maps of central carbon metabolism and light harvesting reactions over a 24 hour period with 12 hour light 12 hour dark conditions. Labels above the flux maps give time since the onset of light, at t = 12 hours the light turns off. The thickness of the arrows indicates the amount of flux in mmol/gDW·hr through the given reaction, and the color indicates inclusion – black lines carry significant flux, gray lines do not, and red lines are not in the model of this organism. The model includes compartmentalization of reactions as shown in the figure: orange represents the mitochondria and green represents the plastid; dotted lines indicate transport flux between these different compartments. It is clear from these select time points that the cell experiences dramatic shifts in carbon and light reactions during the day.

### Modeling Mutant Phenotypes

The model is capable of predicting phenotype based solely on genomic and transcriptomic data. As a validation, we simulated the phenotype of a starchless mutant (*sta6*) (47) in diurnal light both with acetate and on minimal media and a wild type strain in the same conditions (Figure 3A). Based on our simulations, the wild type strain can grow in diurnal light in both conditions, though as expected, in the presence of a reduced carbon source (acetate), growth rate is higher. The *sta6* mutant however, is only capable of growth in the presence of acetate; this is the expected result as a cell that is unable to store starch during the day cannot sustain itself at night and agrees with published data by Davey et al. (3). This also highlights the power of our modeling approach because a normal flux balance analysis (FBA) model would not predict that this gene is essential. Due to the way FBA is formulated, it can only model steady state conditions, not the transient changes that occur during the switch back and forth. This can result in knockouts that are not fatal under steady state becoming so, such as Cre02.g145800.t1.2, a gene that encodes for mitochondrial malate dehydrogenase. As might be expected from a cell that heavily relies upon shuttling carbon to the mitochondrial ATP synthase, removing the cell’s ability to do so efficiently results in long-term failure to thrive. There are also some knockouts that are predicted to overproduce in a circadian environment, despite being fatal in a steady state one – but these are universally the result of the dynamic’s model changing biomass constraints. Because the model does not strictly constrain the ratios of biomass components, the model can suggest that some knockouts are feasible despite their inability to produce biomass components; a good example of this is Cre17.g728950.t1.1, which controls production of flavin adenine dinucleotide (FAD). This redox coenzyme is only required in small amounts in the biomass equation, but is large and therefore metabolically costly to produce. Mutants that cannot make it are predicted to grow faster, but this result is unlikely to be experimentally borne out. A full summary of these knockouts is available in Supplemental File 19.

**Figure 3.**
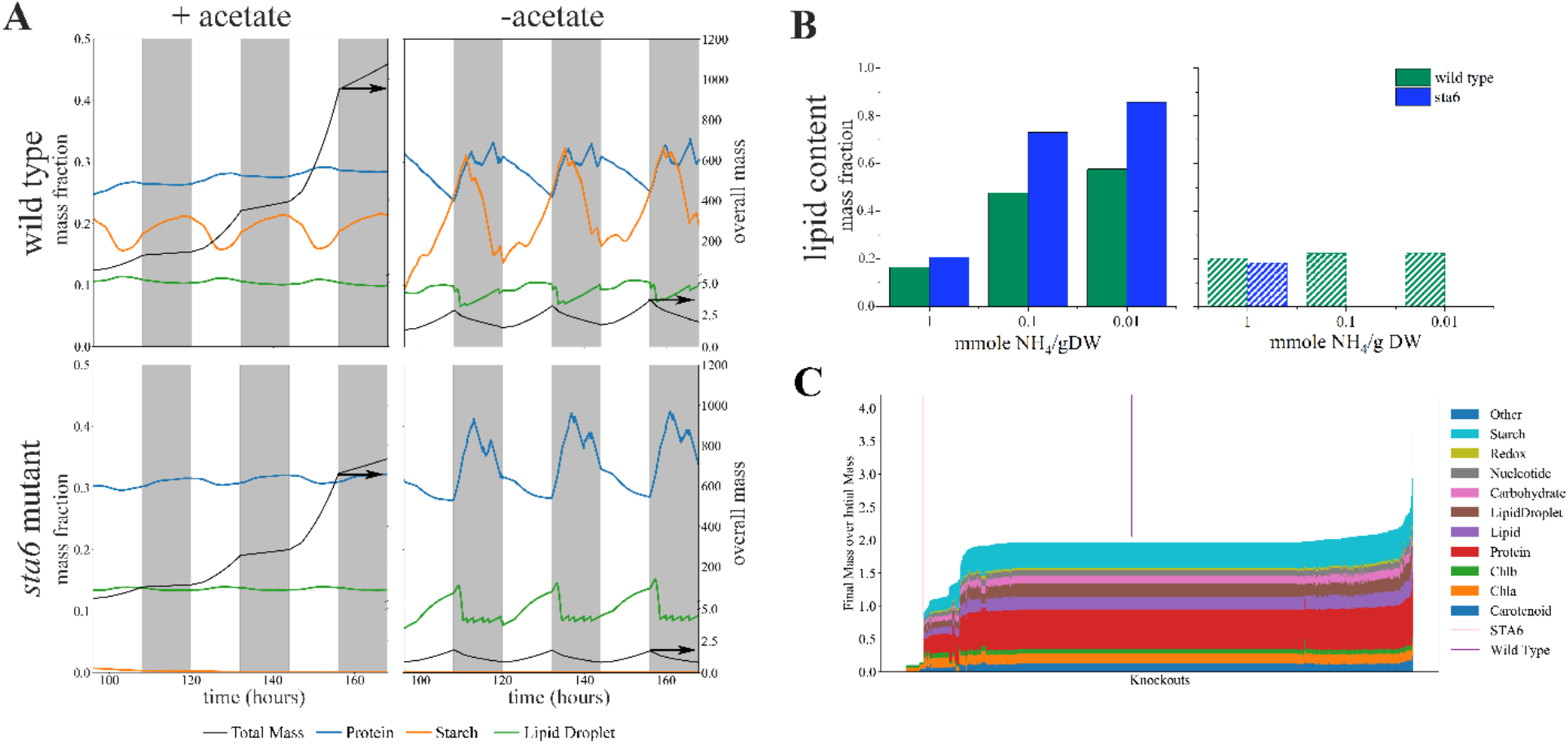
Predicting phenotype from genotype. A) Growth and biomass composition simulations for WT and *sta6* in diurnal light (light indicated in white and night indicated in grey) with and without an organic carbon source. In the presence of acetate, both strains grow well, with the wild type strain outperforming the *sta6* mutant (overall biomass in black line). Without acetate (only CO_2_), the model predicts much slower growth for the WT strain and no growth for the *sta6*. B) Predicted lipid content for WT and *sta6* strains in constant light (solid bars) and in diurnal light (lined bars). C) Phenotype of individual gene knockouts for every gene in the metabolic model. Each vertical line in the graph represents a single gene knockout. The parent strain (WT) doubles once in 24 hours and thus its mass ratio is 2. The majority of gene knockouts have the same final biomass as the WT, but there are many that result in severe defects in growth (on left side of the graph) and several that result in higher biomass.

Our model can predict not only growth and metabolic fluxes, but also changes in biomass composition as well. This feature is incredibly important as most efforts to engineer algae for biofuels have focused on the accumulation of lipids or fatty acids and our model is capable of predicting the effect of environmental or genetic changes on biomass. We used the model to predict the effect of nitrogen limitation on the accumulation of lipids in the wild type and *sta6* mutant in both continuous light and diurnal light (Figure 3B). In continuous light, both strains accumulate more lipids with decreasing nitrogen in the medium and we see the phenotype that has been reported in a number of papers on *sta6* (27, 47–50): that it is capable of accumulating more lipid than the wild type. When we repeated the simulations for growth in diurnal light, the wild type strain loses its ability to respond to nitrogen stress with the accumulation of lipids, it has almost the same lipid content regardless of the nitrogen limitation imposed on the model. As we described earlier, the *sta6* mutant is incapable of growth in diurnal light and this is repeated (and likely exacerbated) with nitrogen limitation. This feature of the model lets us evaluate how external or internal changes in the cell impact biomass composition.

### Genotype to Phenotype Prediction

The model was then used to predict phenotypes of other genetic knockouts (see Figure 3C). A key parameter for economical production of algal based biofuels is productivity; for growth associated products, higher growth rates lead to higher productivities. Therefore, we first focused on the impact these gene knockouts had on growth. Our simulations are based on experimental data that was specifically designed to have one cell doubling in a 24-hour period; therefore, any knockouts that impair growth have a biomass less than 2 and those that result in higher biomass have biomass larger than 2 (Figure 4). Not surprisingly, many of the underperforming mutants are knockouts of critical enzymes involved in pigment biosynthesis, chlorophyll biosynthesis and light harvesting (see supplemental Table S1; supplemental file 20). Table S1 provides more detail for all mutants with a growth defect of 25% or more; of the 146 knockouts in the table, 23 encode enzymes which are part of the GreenCut2 (51), a functional classification of enzymes specific to the plant lineage. We investigating the fuctionality of these genes with Phtyozome (52), and then compared these mutants to the genomewide knockout library constructed and tested for growth rate in minimal medium photoautotrophically by Li et al.; given the major growth defects predicted by the model, it is not surprising that most were not found (53). Of the 1355 genes in the model, only 39 were predicted to result in a 10% or higher increase in biomass (see supplemental Table S2, supplemental file 21). The simulation predicts higher growth in diurnal conditions, but closer inspection of the list can throw out several possibilities prior to constructing the mutant. For example, *nic2, nic7* and *nic13* are auxotrophic for nicotinamide (54) and two strains are deficient in genes encoding enzymes present in the photorespiratory pathway (*GYD1* and *AGT2*), thus requiring high CO_2_ for growth and one strain is lacking a key glyoxylate shunt enzyme (*MAS1*), which is a known carbon conserving mechanism. The other mutants predicted to have better growth have not yet been characterized experimentally for growth in diurnal light.

**Figure 4.**
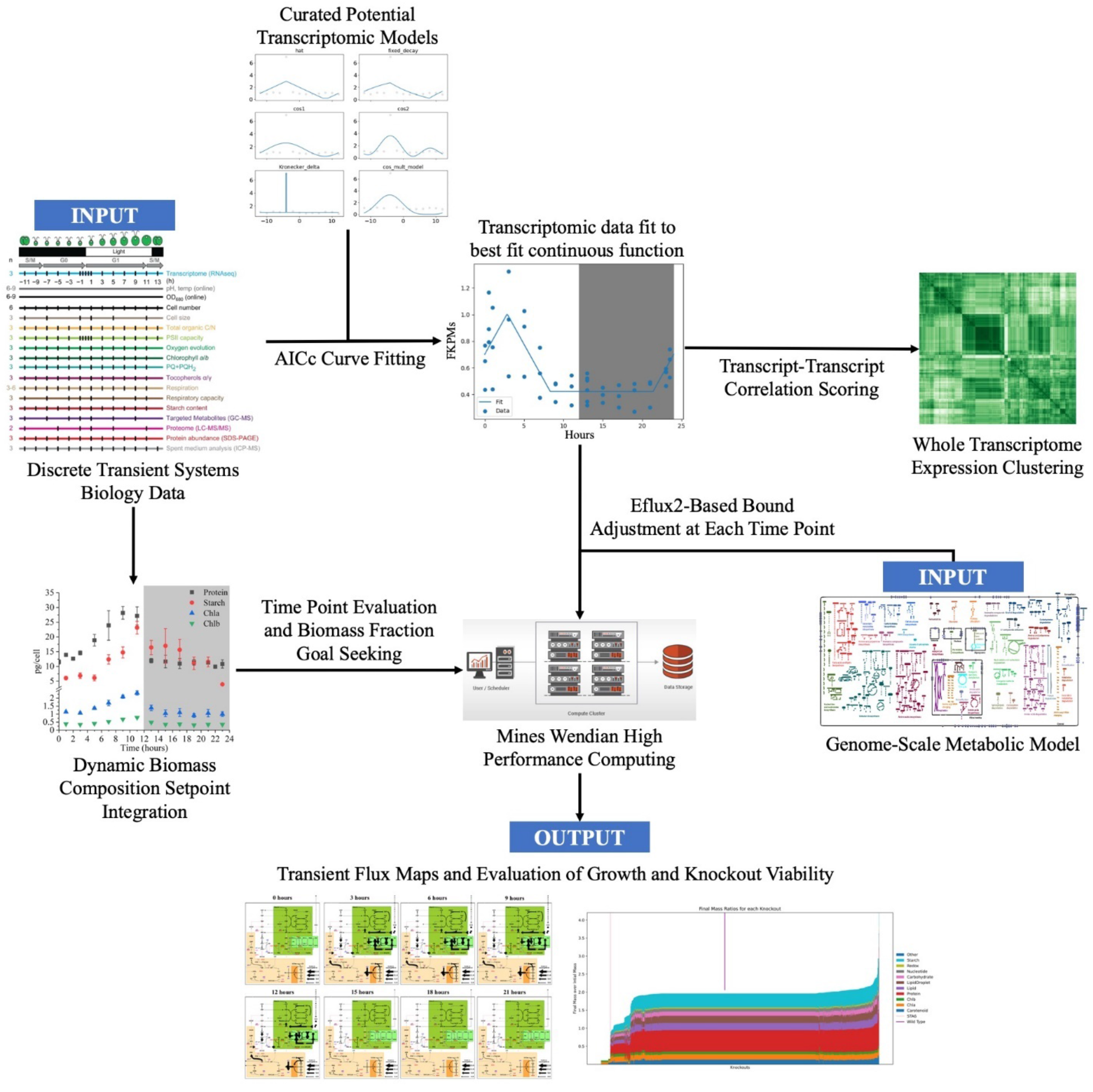
The data pipeline used to create a dynamic metabolic model and a set of correlations. This can be utilized for any algal cell that has these data sets. Transcriptomic correlations aid in the discovery of new interactions between genes; the mechanistic metabolic model enables for the testing of knockouts and media adjustments under dynamic circadian conditions.

## CONCLUSION

While steady state metabolic modeling is an important tool for biological engineering, photoautotrophic species exhibit strong circadian rhythms that are evident both it the cyclic changes in transcript abundance and biomass composition. These transient changes require a modeling approach that allows for the dynamic changes occurring in the cell. Here, we leverage the ease of implementation of dynamic FBA models with gene expression and biomass composition data to develop a model that is capable of predicting phenotype in diurnal light. This model simulates growth in both day and night continuously, accounting for catabolism, changing objective functions, and transcriptomic constraints. We have shown its ability to accurately predict phenotype from genotype, making it a useful tool both for physiological interrogation of biology and metabolic engineering efforts. That predictive power, in combination with the inherently parallelizable nature of *in silico* modeling, means that it can be used to quickly assess potential bioengineering approaches for feasibility, as demonstrated by individual simulations of a broad swath of single knockout mutants.

## MATERIALS AND METHODS

### Informational Pipeline

In order to utilize a large dataset of circadian transcriptomic expression patterns, we built a pipeline for mass-fitting simple expression curves to every single individual transcript. These curves are simple enough to reduce overfitting, but also have enough parameters to preserve critical cyclical information. Once these curves were fit, we used them in two ways. The first was a clustering approach that enabled us to compare how every single transcript’s expression pattern compare with every other transcript within the cell. This produced a heatmap where several clear correlations could be identified, including the collection of genes with a strong response to sudden light onset. The second was the integration of these curves into a genome-scale metabolic model; by querying the transcriptomic curves at every time point, the boundaries of the metabolic model could be dynamically adjusted over the course of the day (Figure 4).

### Fitting Discrete Transcriptomic Data with Continuous Functions

To build this data-driven model of *Chlamydomonas reinhardtii*, we used published transcriptomic data from cells grown in 12:12 day night cycles (1). This data set provided two hour resolution (or less) of changes in gene expression across the entire genome of *Chlamydomonas*. To use this data as constraints for a model with smaller time steps, we first needed to fit the discrete data points with continuous functions. After a quick visual inspection of the data, we chose 8 different types of transcript expression models that would fit the different expression profiles well: One term cosine model, two term cosine model, hat model, fixed decay model, variable decay model, constant model, Kronkecker delta model, and cosine multiplicative model (see supplemental figure S1). For every gene in the transcriptome, we first estimated the parameters for each of the 8 proposed expression profiles which provided the best fit (see supplemental methods). Then, we used the corrected Akaike Information Criterion (AICc) (55, 56) to identify the profile that provided the best fit with the least loss of information.

The parameter estimation for each transcript expression model was performed with scipy.optimize.curve_fit. As there was no guarantee of global optima, the initial conditions for the fitting protocol were randomly perturbated and re-fit 100 times, with the lowest value for the AICc fit being selected for that particular model. Then, all the transcript expression models were compared against each other on the basis of AICc; the lowest overall AICc model was assumed to be the best model for this specific transcript. This protocol was repeated for every single transcript, thereby transforming thousands of sets of discreet data points into a collection of curves that represented the cell’s circadian rhythms.

### Clustering Genes by Expression Profile

The curves assigned to specific transcripts can also be used to evaluate the similarity between each transcript. This is typically performed in two ways: score and integral similarity. As described below, score is based off of vectorizations of the parameters for each curves – by calculating the angle and distance between two fits, a quick measurement of curve similarity can be assessed. Integral similarity between curves is found by sampling the best fit curve for each transcript at small intervals, linearly mapping the sampled values such that the minimum sampled value is zero and the maximum sampled value is one in order to normalize each, and finally comparing the differences between the normalized curves at each sample point. Those with a high integral of difference are very dissimilar curves, while those with a very small integral have highly similar expression shapes (though they may have large differences in expression magnitudes). By calculating these measurements for every possible pair of transcriptomic interactions and clustering the results, we generate a heatmap that summarizes the relationships between the circadian expressions of every gene. This map can be built for either method of comparison, either score or integral; the clustering in this paper was performed with the integral method.

### Scoring

The score method is faster than the integral method at examining correlations between transcripts, and provides a good “first look” at potential clusters. However, it is also much more abstracted than the integral method. By vectorizing the parameters of the curves associated with each transcript, we can compare two vectors with relatively few calculations. Minimizing calculations is critical, as the number of comparisons scales as O(n^2^).

In order to vectorized a given expression curve, we begin by calculating the weight of the transcript expression models associated with the curve. A model’s weight is calculated by comparing the AICc to the AICc’s of all models summed together in the following fashion, where Δ_i_ is the difference between the AICc of model *i* and the lowest AICc within the set of transcript expression models associated with the curve (here represented by *R*):

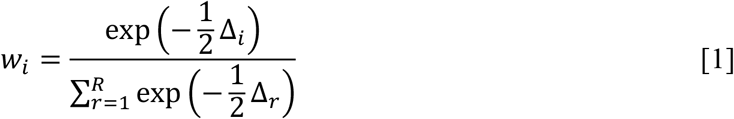

Weight is roughly conceptually equivalent to the probability that a specific model is the best of all given models at fitting the data. Repeating the above equation for all models within the set *R* shows that the weights of all model fits for a particular set of data must sum to 1 (57). When we multiply the weight of a given transcript expression model by the parameter values that provide the best fit for that same model, we transform the fit into a vector that represents both the fit’s parameters and the utility of said fit; repeating this calculation for every model and concatenating all the resulting vectors into one gives us a vector that encapsulates all of the transcript expression models simultaneously.

These vectors enable inexpensive comparison between different transcripts. Because vector operations are, computationally speaking relatively cheap, we can quickly run a number of calculations for each vector comparison. We first calculate the distance between each pair of vectors, using the Euclidian norm. The distance between a pair of vectors can indicate the similarity of fit parameters – even if two curves heavily favor the same model, the difference in parameters will result in a high distance. However, two curves that weight different models strongly may still have a small Euclidian distance between them. To account for this, we also calculate the angle between the vectors. If one transcript heavily weights a particular model while another places a significant weight on another, it is likely that the shape of the transcript expression is quite different. When the angle is calculated between them, it will reflect that information and approach orthogonality. We then combine these pieces of information in a multiplicative fashion, so that if either distance or angle indicates dissimilarity, the score will reflect that fact. While the cosine of the angle is already conveniently normalized between −1 and 1, it’s desirable to also map the score onto a similar space. To do so, we generated a distance value:

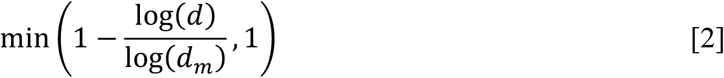

where *d* is the distance between the vectors, and *d_m_* is the maximum distance for all gene ID vector pairs. This is neatly locked between 0, when the distance of the pair is the maximum distance in the entire data set, and 1, when the distance is so close as to be negligible. Moreover, the log operations shift the sensitivity of the curve so that most of the 0-1 range of distance values are mapped onto relatively small distance values, thereby highlight small differences in distance even when maximum distance values are large. Multiplying the cosine of the angle by the distance value gives the score expression:

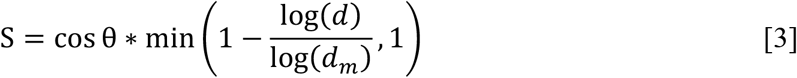

It is immediately clear that the only way to approach the maximum score of 1 is to have vectors that are closely aligned and of similar magnitudes. Should two vectors not be closely aligned, they cannot approach 1, regardless of how distant they are from each other – and should two vectors have near-identical direction but different magnitudes, their score will also be low.

### Image construction

For every clustering map, the dataframe containing the relevant data (either score or integral values for each pair) is loaded. As this file is generally around 6GB, it’s infeasible to generate the entire map at once. Instead, a set of linkages is created using scipy’s cluster.hierarchy.linkage with the ward map. The linkages are then used to produce a truncated dendrogram with a set number of leaves. Each leaf is then processed into a non-square heatmap, where every pixel represents one pair-wise comparison between transcripts. The maps are made iteratively, with each being cleared out of the memory before the next is added. Once all the maps are complete, the dataframe is no longer required and can be cleared out as well, leaving space for each map to be loaded back in and stitched together into a final square map. For the weight sidebar, each model weight is included as a section of a stacked bar graph, which is saved separately and then added to the square map. Additional information can be added if required, such as the highlighting of specific genes.

### Using transcriptomic data to constrain solution space with E-Flux2

Simulating a cell with a constraint-based approach requires constraints, which ideally reflect actual physiological or environmental phenomena. Transcripts can be integrated into these models in order to inform these constraints, using approaches like the E-Flux2 methodology (58); this method multiplies the level of transcriptomic expression, normalized to a fraction of the maximum, by the maximum allowable flux for all reactions, and sets the result as the maximum allowable flux for that specific transcript. In this simulation, the maximum allowable flux through any reaction was set by matching the maximum light uptake through the transcriptomic constraints to the maximum amount of light in the environment.

We then extended this method by dynamically shifting the constraints of all reactions with transcriptomic linkages at every time point. At each time point, each curve was interrogated and normalized against the maximum for that curve over the day, and the bounds lowered to that normalized fraction. When reactions involved more than one gene, the bound for each was calculated and them combined according to the structure of the gene-reaction rule itself (AND relationships took the minimum of the involved fluxes, while OR relationships summed together the fluxes). This is depicted mathematically below; Equation 4 defines the normalization function, Equations 5 and 6 show the equations for AND and OR relationships, respectively, while Equations 7, 8, and 9 show an example.

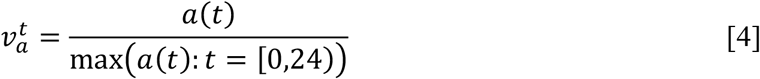

Equation 4’s resulting value, 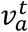, represents the normalized flux fraction for a given gene *a* at time *t*, where *a*(*t*) represents the value of the best-fit curve for gene *a* at time *t*.

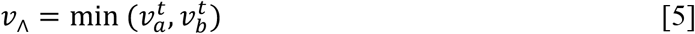

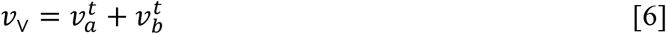

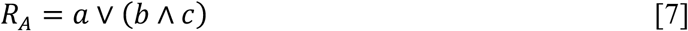

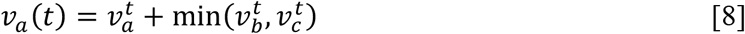

Equation 7 is a mathematical representation of the gene-reaction rule *a* OR (*b* AND *c*); Equation 8 is the corresponding representation of how to calculate the flux fraction for the same rule.

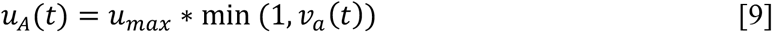

Finally, the flux fraction is multiplied by the maximum reaction flux of the system to calculate the bound for the reaction *A* at time *t*. This value is used for the upper bound of the reaction. If the reaction is reversible, the negative of it is used for the lower as well. Otherwise, the lower bound of the reaction is zero. The maximum reaction flux of the system was calculated such that the maximum rate of light uptake over the course of the day matched the light availability of the system. This approach is feasible, despite the discreet nature of the transcriptomic data set, because the process of fitting the curves to the point creates a continuous representation of the cell’s transcriptome that can be queried at any arbitrary time.

### Decoupling Biomass and Transient Objectives

A typical constraint based model formulation uses a static biomass formation equation to represent metabolites being used for the creation of biomass. Such a fixed equation is inappropriate for a circadian model, both because the composition of the cell changes over the diel cycle and because the cell’s capability to make individual components of the broader equation may change over time. In order to accommodate for this difference, we split the cell’s biomass equation into sub-groups and laid a control system over the top of it, such that the model would always attempt to aim for a long-term stable solution that approached the known biomass composition at different times of the day. The decoupled biomass equation consists of 10 components: Carotenoid, Chlorophyll A, Chlorophyll B, Protein, Lipid, Lipid Droplet, Carbohydrate, Nucleotide, Redox, and Starch; details of these are in supplemental file 22. This division of biomass allows the model to produce different components of biomass when it can, without compromising the cell’s ability to grow if a specific biomass metabolite cannot be made.

In order to properly treat lipid production, we separated the lipid components of biomass into both a general lipid component and a lipid droplet component. The lipid droplet contained all triacylglycerides in the original biomass equation that had tails measured in Wang, et al., while the general lipid class contained all other lipids (2). This separation of lipids allows the lipid droplet fraction to rise and fall independently of the other lipid components. This behavior is pertinent for modeling lipid fraction under nitrogen stress, as the general lipid component contains lipids that contain nitrogen, and therefore will be constrained in times of low nitrogen availability.

### Setting catabolism and anabolism rates

After the model is constrained to a timepoint, the production possibilities need to be evaluated. Critically, this means that the production space for biomass components must be defined. In this modeling approach, the biomass equation of the original FBA model – iCre1355 (40) in this implementation, though this approach is not limited to only this model – is split into components, so that not all biologically relevant metabolites need to be available in the correct ratios for any of them to be consumed. For every timepoint in the diel cycle, the production of each component is tested: first, for production from just the exchange reactions available, and then for production from consumption of each of the other biomass components.

Once the timepoints of the daily cycle have been evaluated, the model begins to run. At each time point, the model is constrained according to the transcriptomic curves, and the production goals and consumption preferences are set by the error terms for the components. These error terms are calculated using a pair of PI control systems – one for the production terms, and one for the consumption terms – thereby abstracting the complex cellular regulatory networks that govern anabolic and catabolic processes into a simpler parameter space. After the production and consumption error terms for a given time point have been calculated, the model tries to produce the non-growth associated maintenance ATP (NGAM) with the media – if it cannot, it will then consume one or more components of biomass to stay alive.

Component consumption is prioritized by the consumption error terms – those with the highest positive error (i.e. the component in question is likely above its setpoint and has been for some time) are consumed first. Once the cell has made NGAM, it evaluates production potential. If the media can be successfully consumed (i.e. it can utilize light to produce NGAM), or if any components with positive degradation error remain after making ATP, the cell will attempt to produce components from either the media or the positive error components, respectively. The production possibilities are precalculated for each time point and possible energy source, either media or specific components. In the event that metabolites in a component cannot be catabolized by the cell, they are assumed to be dumped out of the cell entirely, resulting in an overall mass and energy loss. The cell uses the production error terms and the production possibilities to calculate which components are produced at what ratios. These components and ratios form the initial objective function for the cell’s simulation step.

During the production phase, the cell attempts to produce as much of the initial objective as possible. Once it reaches a limitation, the model fixes the fluxes, then checks which component is limiting production and attempts to make more of the non-limiting desired ones. This iterative expansion of the production window continues until there are no more desired components, or – more commonly – the constraints of the cell limit overall production. Once components have been consumed and produced, the rates of production and consumption are multiplied by the total cell mass and used to update the amounts of each component.

### Universal Single-Knockout Evaluation

In order to demonstrate the utility of this modeling approach, we conducted a survey of all gene knockouts within the model. Using high performance computing (HPC), we were able to simultaneously evaluate large numbers of genes. To simulate each gene knockout, the transcriptomic fits for the gene in question were set to zero, thereby simulating a complete knockout. The knockout mutants were simulated for 168 hours, a full week, at light and non-growth-associated maintenance ATP light levels that produced approximately one doubling in the wild type cell over the length of the simulation. For every run, the full biomass composition was recorded at the end of the simulation, and the impacts of each gene on the overall growth and biomass composition can be clearly seen.

## ACKNOWLEDGEMENTS

We would like to thank Dr. Sabeeha Merchant and Dr. Daniela Strenkert for their valuable discussion during the development of the model. This research was supported by the DOE Office of Science, Office of Biological and Environmental Research (BER), grant no. DE-SC0019171

