## Supplemental File 1 for "Rhythm of The Night (and Day): Predictive metabolic modeling of circadian growth in *Chlamydomonas*"

**Additional Supplemental Files**

- Supplemental_File_2.xlsx – an excel sheet of FPKM data and functionality for the *RACK1* clustering neighborhood.
- Supplemental_File_3.xlsx – an excel sheet of FPKM data and functionality for the *LHSCR3.1* clustering neighborhood.
- Supplemental_File_4.xlsx – an excel sheet of FPKM data and functionality for the *RBCS1* clustering neighborhood.
- Supplemental_File_5.xlsx – an excel sheet of FPKM data and functionality for the *FAP85* clustering neighborhood.
- Supplemental_File_6.mp4 – a movie of central metabolic flux distributions.
- Supplemental_File_7.png through Supplemental_File_18.png – plots of the bounds and fluxes through various photosynthetic reactions over the course of a simulated day.
  - 7 – TPIh
  - 8 – TKT2h
  - 9 – TKT1h
  - 10 – TAh
  - 11 – RBPCh
  - 12 – PSIred
  - 13 – PSIIred
  - 14 – PGKh
  - 15 – FBA3hi
  - 16 – Photon Exchange
  - 17 – CEF
  - 18 – CBFC
- Supplemental_File_19.xlsx – an excel sheet of all knockout effects.
- Supplemental_File_20.xlsx – an excel sheet of all underperformers, summarized in table S1 below.
- Supplemental_File_21.xlsx – an excel sheet of all overperformers, summarized in table S2 below.
- Supplemental_File_22.xlsx – an excel sheet of biomass components.

**Github**

Additional result files and source code are available at https://github.com/metcalex/Transcriptomic_Circadian_Modeling_Supplemental

**Model Updates**

In order to utilize the iCre1355 model for this approach, we had to modify the way biomass and excretion were handled. Every biomass component was assigned a specific classification, as shown in Supplemental File 2. This split the biomass equation into ten sub-equations, where each sub-equation preserves the relative ratio among individual metabolites within the same classification. The sub-equations were then normalized by mass, so that 1 mmol of flux through a sub-equation produced 1 gram of mass imbalance. These normalized equations were added to the model.

We also added excretion equations for every single component within the biomass equation, so that the cell could dump components it was unable to catabolize. This meant that any biomass classification could be consumed for energy, as the steady-state limitation no longer applied; even if no degradation pathway exists for one metabolite in the sub-equation, it can be excreted while the rest are degraded for energy and/or carbon.

**SUPPLEMENTAL FIGURES**

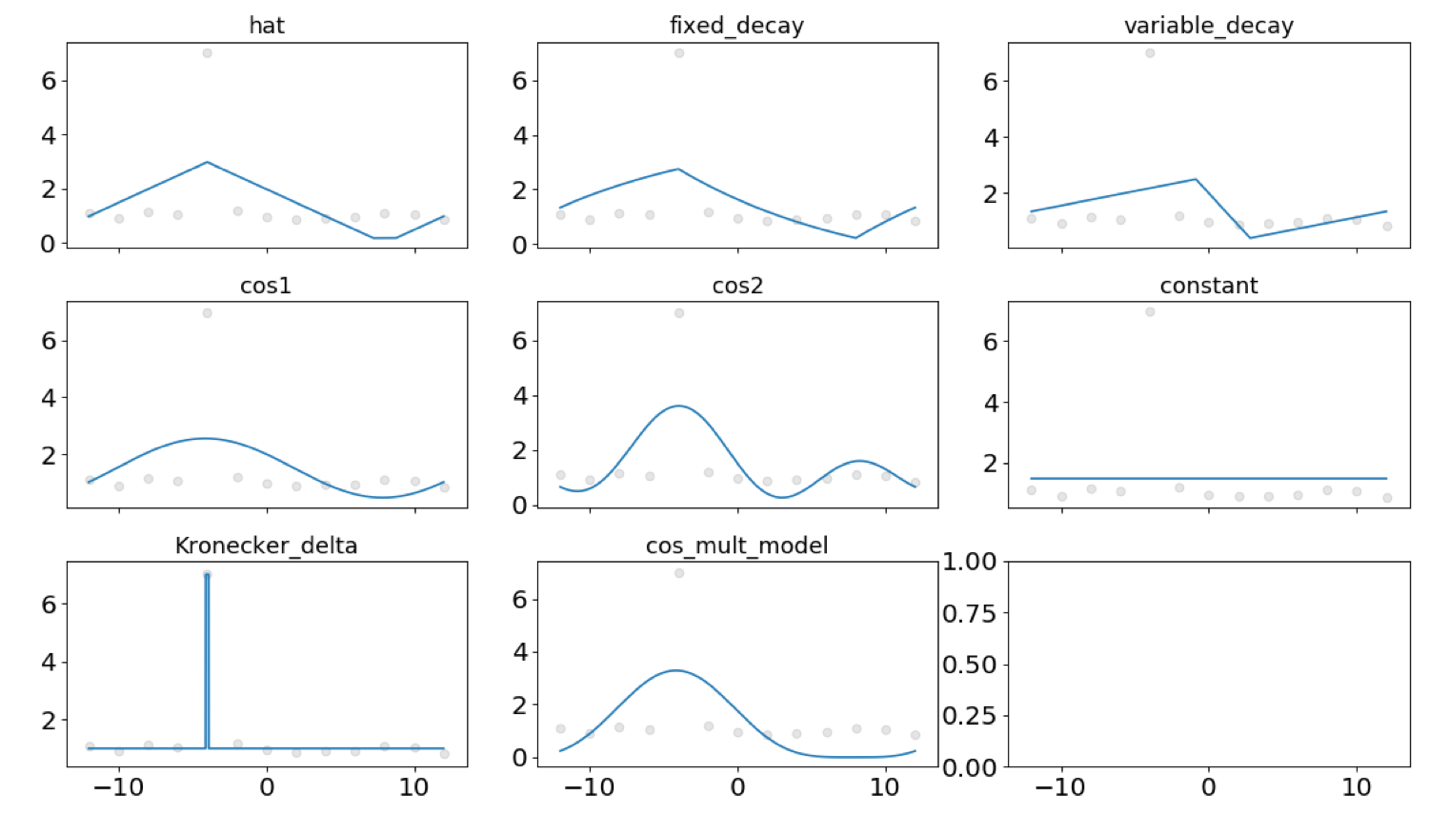

**Figure S1.** **Examples of the different models used to describe transcriptomic data.**

**SUPPLEMENTAL TABLES**

**Table S1. Predicted growth of mutants which underperform the parent strain (wild type has mass of 2) by >25% sorted by mass.**

| **Transcript ID** | **Gene Name** | **Functionality** | **GreenCut2?** | **Li et al. Underperformer?** | **Predicted Mass of Knockout Strain** |
| --- | --- | --- | --- | --- | --- |
| Cre16.g687900.t1.2 | LHCA7 | Light-harvesting protein of photosystem I | N | Y | 0.104 |
| Cre10.g452050.t1.2 | LHCA4 | light-harvesting protein of photosystem I | Y | N | 0.104 |
| Cre06.g272650.t1.2 | LHCA8 | Light-harvesting protein of photosystem I | N | N | 0.104 |
| Cre11.g467689.t1.1 | PETC | Chloroplast cytochrome b6f Rieske iron-sulfur center subunit | Y | Y | 0.104 |
| Cre02.g082500.t1.1 | PSAN | Photosystem I reaction center subunit N | N | N | 0.104 |
| Cre12.g486300.t1.2 | PSAL | Photosystem I reaction center subunit L | Y | Y | 0.104 |
| Cre05.g238332.t1.1 | PSAD | Photosystem I reaction center subunit II, 20 kDa | Y | Y | 0.104 |
| Cre10.g420350.t1.2 | PSAE | Photosystem I reaction center subunit IV, 8.1 kDa | Y | Y | 0.104 |
| Cre09.g412100.t1.2 | PSAF | Photosystem I reaction center subunit III | Y | N | 0.104 |
| Cre17.g724300.t1.2 | PSAK | Photosystem I reaction center subunit K | Y | N | 0.104 |
| ChreCp045 | petG | Cytochrome b6-f complex subunit 5 | N | N | 0.104 |
| Cre12.g558900.t1.2 | PETO | Regulator of photosynthetic cyclic electron flow | N | N | 0.104 |
| Cre07.g340200.t1.1 | PGRL1 | Proton-gradient related-like | Y | N | 0.104 |
| Cre14.g626700.t1.2 | PETF | Chloroplast ferredoxin | Y | N | 0.104 |
| ChreCp001 | petA | Cytochrome f | N | N | 0.104 |
| Cre12.g546150.t1.2 | PETM | Chloroplast cytochrome b6f PetM subunit | N | Y | 0.104 |
| Cre11.g476750.t1.2 | FNR1 | Ferredoxin-NADP reductase, chloroplast | Y | N | 0.104 |
| Cre12.g550850.t1.2 | PSBP1 | Photosystem II Oxygen Evolution Enhancer protein 2 | Y | Y | 0.104 |
| Cre10.g440450.t1.2 | PSB28 | Photosystem II associated subunit 28 | Y | N | 0.104 |
| ChreCp001 | psbF | Cytochrome b559 subunit beta | N | N | 0.104 |
| ChreCp051 | psbI | Photosystem II reaction center protein I | N | N | 0.104 |
| ChreCp029 | psbH | Photosystem II reaction center protein H | N | N | 0.104 |
| Cre08.g372450.t1.2 | PSBQ | Photosystem II Oxygen Evolution Enhancer protein 3 | Y | N | 0.104 |
| ChreCp027 | psbZ | Photosystem II reaction center protein Z | N | N | 0.104 |
| Cre06.g273700.t1.2 | HCF136 | Photosystem II stability/assembly factor HCF136 | N | Y | 0.104 |
| Cre16.g673650.t1.1 | LHCB5 | Light-harvesting minor chlorophyll a/b binding protein of photosystem II | Y | N | 0.104 |
| Cre05.g243800.t1.2 | PSB27 | conserved expressed protein involved in PSII biogenesis | N | Y | 0.104 |
| ChreCp021 | psbA | Photosystem II protein D1 | N | N | 0.104 |
| ChreCp066 | psbC | Photosystem II CP43 reaction center protein | N | N | 0.104 |
| Cre08.g362900.t1.1 | PSBP4 | PsbP-like protein anchored in thylakoid lumen | Y | N | 0.104 |
| Cre07.g328200.t1.2 | PSBP6 | PsbP-like protein of thylakoid lumen | Y | N | 0.104 |
| Cre01.g016600.t1.2 | PSBS1 | chloroplast Photosystem II-associated 22 kDa protein | Y | N | 0.104 |
| ChreCp031 | psbT | Photosystem II reaction center protein T | N | N | 0.104 |
| Cre10.g452100.t1.1 | PSBY | ycf32-related polyprotein of photosystem II | N | N | 0.104 |
| Cre07.g328250.t1.2 |  | Pumilio-family RNA binding protein | N | N | 0.104 |
| Cre03.g185550.t1.2 | SEBP1 | Sedoheptulose-1,7-bisphosphatase | N | Y | 0.104 |
| Cre01.g029300.t1.2 | TPIC1 | Triose phosphate isomerase | N | N | 0.137 |
| Cre11.g467569.t1.1 | ATPD | Chloroplast ATP synthase delta chain | N | N | 0.15 |
| ChreCp023 | atpE | ATP synthase epsilon chain, chloroplastic | N | N | 0.15 |
| ChreCp054 | atpF | ATP synthase subunit b, chloroplastic | N | N | 0.15 |
| Cre11.g481450.t1.2 | ATPG | Chloroplast ATP synthase subunit II | Y | N | 0.15 |
| ChreCp062 | atpI | ATP synthase subunit a, chloroplastic | N | N | 0.15 |
| ChreCp050 | atpA | ATP synthase subunit alpha, chloroplastic | N | N | 0.15 |
| ChreCp058 | atpB | ATP synthase subunit beta, chloroplastic | N | N | 0.15 |
| Cre06.g259900.t1.2 | ATPC | Chloroplast ATP synthase gamma chain | N | N | 0.15 |
| ChreCp053 | atpH | ATP synthase subunit c, chloroplastic | N | N | 0.15 |
| Cre03.g188250.t1.2 | STA6 | ADP-glucose pyrophosphorylase small subunit | N | N | 0.82 |
| Cre06.g286250.t1.2 | MPC1 | Mitochondrial substrate carrier protein | N | N | 0.829 |
| Cre08.g358580.t1.1 | CMP1 | Carbamoyl phosphate synthase, large subunit | N | N | 0.895 |
| Cre02.g145800.t1.2 | MDN3 | NAD-dependent malate dehydrogenase | N | N | 0.919 |
| Cre06.g308500.t1.2 | CMP2 | Carbamoyl phosphate synthase, small subunit | N | N | 0.922 |
| Cre05.g234638.t1.1 |  | Possible Amidophosphoribosyltransferase / Phosphoribosyldiphosphate 5-amidotransferase | N | N | 0.928 |
| Cre06.g260200.t1.2 | MCP18 | Mitochondrial substrate carrier protein | N | N | 0.951 |
| Cre08.g364800.t1.2 |  | Possible Phosphoribosylformylglycinamidine synthase / Phosphoribosylformylglycinamidine synthetase | N | N | 0.957 |
| Cre12.g537581.t1.1 |  | Possible IMP cyclohydrolase / Inosinicase | N | N | 0.957 |
| Cre13.g566000.t1.2 |  | Putative monofunctional formate-tetrahydrofolate ligase | N | N | 0.962 |
| Cre17.g734100.t1.2 |  | Possible Adenylosuccinate synthase / Succinoadenylic kinosynthetase | N | N | 0.965 |
| Cre10.g433600.t1.2 |  | Possible Methylenetetrahydrofolate reductase (NAD(P)H) / N(5,10)-methylenetetrahydrofolate reductase | N | N | 0.968 |
| Cre06.g250902.t1.1 | METH | Cobalamin-independent methionine synthase | N | N | 0.968 |
| Cre07.g318750.t1.2 | PURM | Aminoimidazole Ribonucleotide synthetase | N | N | 0.969 |
| Cre12.g550700.t1.2 |  | Possible Phosphoribosylglycinamide formyltransferase / Glycinamide ribonucleotide transformylase | N | N | 0.998 |
| Cre13.g572200.t1.2 |  | Possible tyrosine-specific transport protein (tyrP) | N | N | 1.043 |
| Cre11.g467770.t1.1 | PGK1 | Phosphoglycerate kinase, chloroplast precursor | N | N | 1.047 |
| Cre08.g368950.t1.2 | DGS1 or DHQS (name change) | 3-dehydroquinate synthase | N | N | 1.073 |
| Cre04.g214150.t1.1 | THI4 | Thiazole biosynthetic enzyme | N | N | 1.08 |
| Cre03.g204250.t1.2 | SAH1 | S-Adenosyl homocysteine hydrolase | N | N | 1.083 |
| Cre06.g250200.t1.2 | METM | S-adenosylmethionine synthetase | N | N | 1.083 |
| Cre03.g160500.t1.2 | TSK1 | Lysyl-tRNA synthetase | N | N | 1.085 |
| Cre03.g193800.t1.1 | TSN1 | Aspartyl-tRNA synthetase | N | N | 1.091 |
| Cre17.g705850.t1.2 | GGK2 | Glutamate 5-kinase | N | N | 1.092 |
| Cre03.g144627.t1.1 | CGS1 | Cystathionine gamma-synthase | N | N | 1.094 |
| Cre02.g082200.t1.1 | HSK1 | Homoserine kinase | N | N | 1.099 |
| Cre17.g726750.t1.2 | SHKA1 | 3-deoxy-D-arabino-heptulosonate 7-phosphate synthetase | N | N | 1.1 |
| Cre01.g050100.t1.2 | DPD1 | Diaminopimelate decarboxylase | N | N | 1.102 |
| Cre10.g434750.t1.2 | AAI1 | Acetohydroxy acid isomeroreductase | N | N | 1.106 |
| Cre08.g380201.t1.1 | SHKD1 | Putative dehydroquinate dehydratase/shikimate:NADP oxidoreductase | N | N | 1.106 |
| Cre16.g672385.t1.1 | HIS5 | Histidinol phosphate aminotransferase | N | N | 1.107 |
| Cre17.g734200.t1.2 | DPA1 | LL-diaminopimelate aminotransferase, putative | Y | N | 1.108 |
| Cre16.g690319.t1.1 |  | probable nucleoside transporter | N | N | 1.109 |
| Cre01.g021251.t1.1 | ARG7 | Argininosuccinate lyase | N | N | 1.11 |
| Cre13.g576650.t1.2 | HIS4 | Bifunctional phosphoribosyl-AMP cyclohydrolase/phosphoribosyl-ATP pyrophosphatase | N | N | 1.11 |
| Cre13.g602350.t1.2 |  | Possible indole-3-glycerol phosphate synthase (trpC) | N | N | 1.111 |
| Cre07.g325400.t1.2 | LEU3 | Isopropylmalate dehydrogenase | N | N | 1.111 |
| Cre06.g279150.t1.2 | TSD2 | Aspartyl-tRNA synthetase | N | N | 1.112 |
| Cre09.g386758.t1.1 | ALS1 | Acetolactate synthase, large subunit | N | N | 1.112 |
| Cre01.g055453.t1.1 | ALS2 | Acetolactate synthase, small subunit | N | N | 1.112 |
| Cre09.g416050.t1.2 | AGS1 | Argininosuccinate synthase | N | N | 1.113 |
| Cre03.g154550.t1.1 | PCR1 | Pyrroline-5-carboxylate reductase | N | N | 1.114 |
| Cre08.g365600.t1.2 |  | Phosphomethylpyrimidine kinase | N | N | 1.114 |
| Cre03.g146527.t1.1 | GSD1 | Glutamic-gamma-semialdehyde dehydrogenase | N | N | 1.117 |
| Cre09.g389689.t1.1 | ASD1 | Aspartate semialdehyde dehydrogenase | N | N | 1.12 |
| Cre05.g240850.t1.2 | THIC | Hydroxymethylpyrimidine phosphate synthase | N | N | 1.121 |
| Cre12.g489700.t1.2 | OTC1 | Ornithine carbamoyltransferase | N | N | 1.123 |
| Cre14.g620300.t1.2 | ASB1 | Anthranilate synthase, beta subunit | N | N | 1.123 |
| Cre06.g278163.t1.1 | ARG9 | Acetylornithine aminotransferase | N | N | 1.124 |
| Cre12.g519000.t1.1 | ASB2 | Phosphoribosylanthranilate isomerase | N | N | 1.124 |
| Cre11.g481500.t1.2 | HIS7 | Imidazole glycerol phosphate synthase | N | N | 1.13 |
| Cre03.g146187.t1.1 |  | Possible N-acetyl-gamma-glutamyl-phosphate reductase / NAGSA dehydrogenase | N | N | 1.131 |
| Cre02.g107300.t1.2 | DPS1 | Dihydrodipicolinate synthase | N | N | 1.132 |
| Cre02.g142352.t1.1 |  | Possible Histidine--tRNA ligase / Histidyl-tRNA synthetase | N | N | 1.136 |
| Cre03.g155200.t1.1 | CHM1 | Chorismate mutase | N | N | 1.137 |
| Cre05.g237400.t1.2 | DAE1 | Diaminopimelate epimerase | N | N | 1.139 |
| Cre02.g077350.t1.2 | HDH1 | Histidinol dehydrogenase | N | N | 1.139 |
| Cre02.g103850.t1.2 | HIS3 | Imidazoleglycerol-phosphate dehydratase | N | N | 1.14 |
| Cre02.g143200.t1.1 | TSA1 | Alanyl-tRNA-synthetase | N | N | 1.141 |
| Cre10.g429150.t1.2 | PRT1 | Anthranilate phosphoribosyltransferase | N | N | 1.141 |
| Cre03.g181300.t1.2 | SHKG1 | 5-enolpyruvylshikimate-3-phosphate (EPSP) synthase (EC 2.5.1.19) | N | N | 1.142 |
| Cre03.g206600.t1.2 | AAD1 | Acetohydroxyacid dehydratase | N | N | 1.143 |
| Cre12.g528700.t1.2 | TRPA1 | Tryptophan synthetase alpha subunit | N | N | 1.144 |
| Cre03.g161400.t1.2 | MAA7 | Tryptophan synthase beta subunit | N | N | 1.144 |
| Cre10.g458050.t1.2 | BCA3 | Branched chain amino acid aminotransferase | N | N | 1.148 |
| Cre03.g143887.t1.1 |  | Arginyl-tRNA Synthetase | N | N | 1.153 |
| Cre06.g261800.t1.2 |  | Prephenate dehydratase | N | N | 1.163 |
| Cre03.g145747.t1.1 |  | Chorismate synthase | N | N | 1.165 |
| Cre01.g004500.t1.2 | LEU1 | Isopropylmalate dehydratase, large subunit | N | N | 1.173 |
| Cre06.g252650.t1.2 | LEU1S | Isopropylmalate dehydratase, small subunit | N | N | 1.173 |
| Cre01.g050950.t1.2 |  | Geranylgeranyl diphosphate reductase / Geranylgeranyl reductase | N | N | 1.284 |
| Cre03.g158000.t1.2 | GSA1 | Glutamate-1-semialdehyde aminotransferase | N | N | 1.293 |
| Cre09.g406050.t1.2 | PYR1 | CTP synthase | N | N | 1.297 |
| Cre17.g715900.t1.2 |  | Similar to Dihydrofolate Reductase-Thymidylate | N | N | 1.304 |
| Cre09.g396300.t1.2 | PPX1 | Protoporphyrinogen oxidase | N | N | 1.308 |
| Cre13.g564650.t1.1 | MRS5 | Magnesium and cobalt transport protein | N | N | 1.311 |
| Cre01.g007737.t1.1 | FBT1 | Folate/pteridine transporter | N | N | 1.319 |
| Cre01.g015350.t1.1 | POR1 | Light-dependent protochlorophyllide reductase | Y | N | 1.32 |
| Cre12.g492950.t1.2 | RIR1 | Ribonucleoside-diphosphate reductase R1 subunit | N | N | 1.32 |
| Cre02.g079700.t1.2 | PYR2 | Aspartate carbamoyltransferase | N | N | 1.321 |
| Cre01.g048950.t1.2 | PYR5 | Uridine 5'- monophosphate synthase | N | N | 1.322 |
| Cre12.g490350.t1.1 | HDS1 | 1-hydroxy-2-methyl-2-(E)-butenyl 4-diphosphate synthase | N | N | 1.335 |
| Cre05.g242000.t1.2 | CHLD | Magnesium chelatase subunit D, chloroplast precursor | Y | Y | 1.337 |
| Cre06.g287750.t1.2 | PYR8 | Dihydroorotase dehydrogenase | N | N | 1.34 |
| Cre01.g042800.t1.2 | DVR1 | 3,8-divinyl protochlorophyllide a 8-vinyl reductase, chloroplast precursor | Y | Y | 1.341 |
| Cre09.g409100.t1.2 | UROS1 | Uroporphyrinogen-III synthase | N | N | 1.344 |
| Cre06.g294750.t1.2 | CHLG | Chlorophyll synthetase | Y | N | 1.347 |
| Cre11.g467550.t1.2 |  | Dihydroorotase/amidohydrolase | N | N | 1.349 |
| Cre06.g272150.t1.2 |  | Possible Protein-disulfide reductase | N | N | 1.351 |
| Cre12.g546050.t1.2 | DXR1 | 1-deoxy-D-xylulose 5-phosphate reductoisomerase, chloroplast precursor | N | N | 1.351 |
| Cre03.g207700.t1.1 | FPS1 | Farnesyl pyrophosphate synthase | N | N | 1.354 |
| Cre16.g669550.t1.2 | METC | Cystathionine beta-lyase | N | N | 1.357 |
| Cre16.g679669.t1.1 | CMS1 | 4-diphosphocytidyl-2C-methyl-D-erythritol synthase, chloroplast precursor | N | N | 1.359 |
| Cre12.g503550.t1.2 | MEC1 | 2-C-methyl-D-erythritol 2,4-cyclodiphosphate synthase | N | N | 1.369 |
| Cre02.g145050.t1.2 | CMK1 | 4-diphosphocytidyl-2-C-methyl-D-erythritol kinase, | N | N | 1.374 |
| Cre08.g372950.t1.2 | IDS1 | 4-hydroxy-3-methylbut-2-enyl diphosphate reductase | N | N | 1.389 |
| Cre02.g091050.t1.2 | ALAD | Delta-aminolevulinic acid dehydratase | N | N | 1.393 |
| Cre07.g342150.t1.2 | HEMA1 | Glutamyl-tRNA reductase | N | N | 1.405 |
| Cre12.g517150.t1.1 | MET16 | Adenylylphosphosulfate reductase | N | N | 1.41 |
| Cre07.g356350.t1.1 | DXS1 | 1-deoxy-D-xylulose 5-phosphate synthase, chloroplast precursor | N | N | 1.412 |

**Table S2. Predicted growth of mutants which overperform the parent strain (wild type has mass of 2) by >10%, sorted by mass.**

| **Transcript ID** | **Gene Name** | **Functionality** | **GreenCut2?** | **Li et al. Underperformer?** | **Predicted Mass of Knockout Strain** |
| --- | --- | --- | --- | --- | --- |
| Cre16.g694850.t1.2 | NGS1 | N-acetylglutamate synthase | N | N | 2.200 |
| Cre17.g735950.t1.2 |  | Possible 2-oxo-4-hydroxy-4-carboxy-5-ureidoimidazoline decarboxylase / OHCU decarboxylase | N | N | 2.201 |
| Cre04.g224150.t1.2 |  | Possible glycerol kinase / Glycerokinase | N | N | 2.202 |
| Cre02.g095137.t1.1 |  | Possible pyruvate synthase / Pyruvic-ferredoxin oxidoreductase | N | N | 2.209 |
| Cre02.g095137.t2.1 |  | Possible pyruvate synthase / Pyruvic-ferredoxin oxidoreductase | N | N | 2.209 |
| Cre09.g413750.t1.1 | AGT2 | Putative aminotransferase | N | Y | 2.216 |
| Cre12.g559950.t1.2 | MFDX | Adrenodoxin-like ferredoxin, mitochondrial | N | N | 2.222 |
| Cre06.g278210.t1.1 | GPM1 | Phosphoglucomutase | N | N | 2.226 |
| Cre17.g698650.t1.1 | GTP1 | Gamma-glutamyl transpeptidase | N | N | 2.240 |
| Cre02.g146050.t1.2 | ATO2 | Acetyl-CoA acyltransferase | N | N | 2.253 |
| Cre02.g092900.t1.2 | GUA1 | GMP synthetase | N | N | 2.269 |
| Cre07.g339554.t1.2 |  | Possible MFS transporter, PAT family, solute carrier family 33 (acetyl-CoA transportor) | N | N | 2.276 |
| Cre01.g022650.t1.2 |  | Possible Beta-ureidopropionase | N | N | 2.301 |
| Cre02.g097000.t1.2 | DHP1 | Dihydropyrimidinase | N | N | 2.312 |
| Cre16.g675650.t1.2 | ALD8 | Aldehyde dehydrogenase | N | N | 2.323 |
| Cre06.g288700.t1.2 | GYD1 | Glycolate dehydrogenase | N | N | 2.342 |
| Cre08.g378150.t1.1 | GLD2 | Glucose-6-phosphate dehydrogenase | N | N | 2.347 |
| Cre10.g435250.t1.1 | NIC13 | NAD+ synthase (glutamine-hydrolyzing) | N | N | 2.364 |
| Cre17.g728950.t1.1 | RFD2 | diaminohydroxyphosphoribosylaminopyrimidine deaminase | Y | N | 2.376 |
| Cre02.g141200.t1.2 | NIC2 | Quinolinate phosphoribosyl transferase | N | N | 2.376 |
| Cre14.g620350.t1.2 | GCH3 | Putative GTP cyclohydrolase | N | N | 2.376 |
| Cre13.g578750.t1.2 | TBA1 | Translation factor for chloroplast psbA mRNA | N | N | 2.393 |
| Cre06.g296600.t1.2 | RFS1 | Riboflavin synthase | N | N | 2.400 |
| Cre17.g708800.t1.1 | GSH2 | Glutathione synthetase | N | N | 2.400 |
| Cre06.g251450.t1.1 | NIC7 | Quinolinate synthetase A | N | N | 2.400 |
| Cre12.g528450.t1.2 | ASO1 | L-aspartate oxidase | N | N | 2.400 |
| Cre02.g088850.t1.2 | RFS2 | Riboflavin synthase | N | N | 2.413 |
| Cre12.g555951.t1.1 | RFD1 | Riboflavin metabolism-associated deaminase | N | N | 2.413 |
| Cre16.g665100.t1.1 | CAV3 | Voltage-gated Ca2+ channel, alpha subunit | N | N | 2.465 |
| Cre03.g144807.t1.1 | MAS1 | Malate synthase | N | N | 2.489 |
| Cre03.g188800.t1.1 |  | Nicotinate phosphoribosyltransferase | N | N | 2.509 |
| Cre12.g500150.t1.1 | ALD5 | Aldehyde dehydrogenase | N | N | 2.676 |
| Cre09.g394658.t1.1 | LCI28 | Low-CO2-induced aldose reductase | N | N | 2.676 |
| Cre17.g722150.t1.2 | PKS3 | Type III polyketide synthase | N | N | 2.934 |
| Cre09.g398289.t1.1 |  | Related to plastidic lysophosphatidic acid acyltransferase (LPAAT) | N | N | 2.936 |
| Cre06.g273250.t1.2 |  | Glycerol-3-phosphate acyltransferase | N | N | 2.936 |
| Cre17.g723650.t1.2 | ATO1 | Acetyl-CoA acyltransferase | N | N | 3.629 |
| Cre06.g308100.t1.2 |  | Possible Enoyl-CoA hydratase 2 / ECH2 | N | N | 4.204 |

**List of abbreviations**

AcCoA: Acetyl- Coenzyme A

AICc: Akaike Information Criterion

AKG: α-ketoglutarate

ATO1: Acetyl-CoA acyltransferase

CoA: Coenzyme A

CIT: citrate

DHAP: dihydroxyacetone phosphate

E4P: D-erythrose-4-phosphate

F6P: fructose 6-phosphate

FAP85: Flagellar associated protein

FBP: fructose biphosphate

FBA: Flux Balance Analysis

FEO: ferredoxin, oxidized

FER: ferredoxin, reduced

FoC: Ferrous cytochrome c

FUM: fumarate

GAP: glyceraldehyde 3-phosphate

G6P: glucose 6-phosphate

3GP: 3-phosphoglycerate

2PG: 2-phosphoglycerate

HPC: High performance computing

ICIT: isocitrate

LHSCR3.1: Light-harvesting complex stress-related protein 3.1, chloroplastic

MAL: malate

NGAM: non-growth associated maintenance

OA: oxaloacetate

PC1+: plastocyanin, reduced

PC2+, plastocyanin, oxidized

PEP: phosphoenolpyruvate

PYR: pyruvate

PQO: plastoquinone, oxidized

PQR: plastoquinone, reduced

QH_2_: Ubiquinone

RACK1: Receptor of activated protein C kinase 1

RBCS1: Ribulose bisphosphate carboxylase small subunit, chloroplastic 1

RNA: Ribonucleic acid

Ru5P: ribulose-5-phosphate

RuBisCo: Ribulose-1,5-bisphosphate carboxylase-oxygenase

SBP: sedoheptulose biphosphate

S7P: sedoheptulose 7-phosphate

SUCC: succinate

SUCCoA: Succinyl-coenzyme A

X5P: xylulose 5-phosphate

Z: photon.
