## Supplementary figures and images for "Rhythm of The Night (and Day): Predictive metabolic modeling of circadian growth in *Chlamydomonas*"

### Supplemental File 7

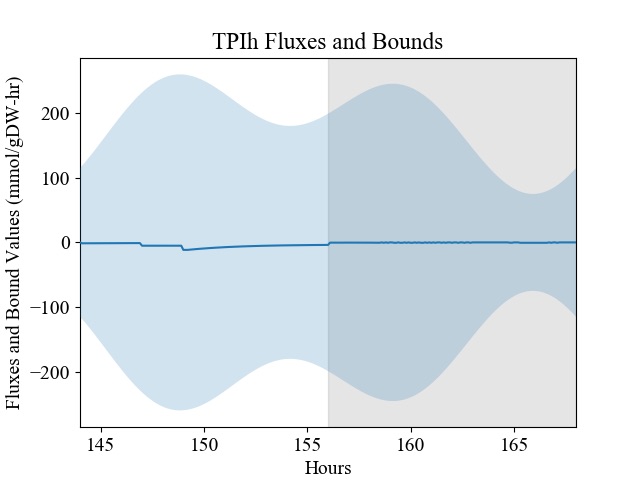

### Supplemental File 8

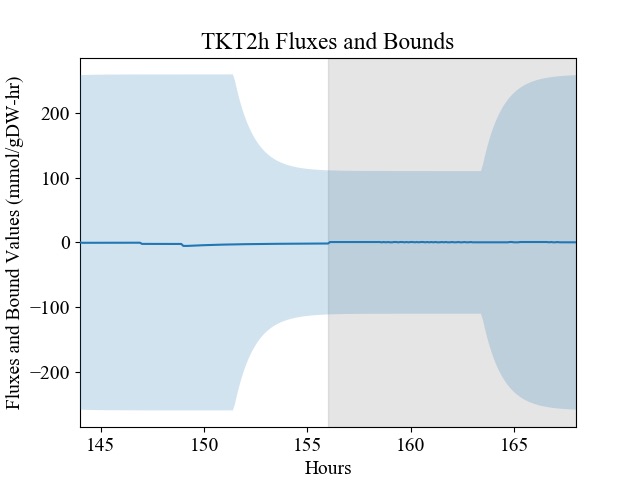

### Supplemental File 9

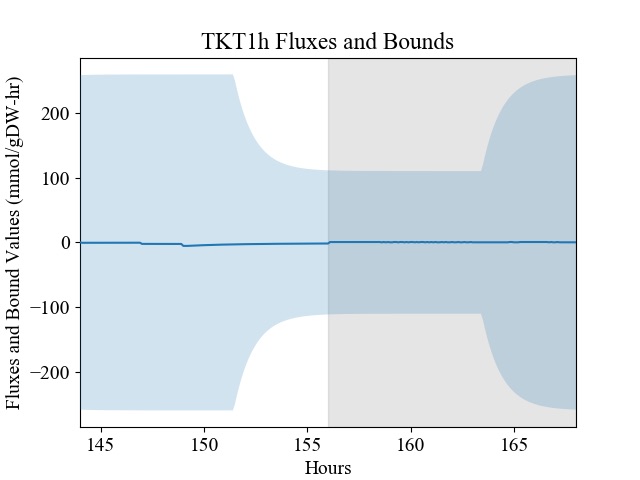

### Supplemental File 10

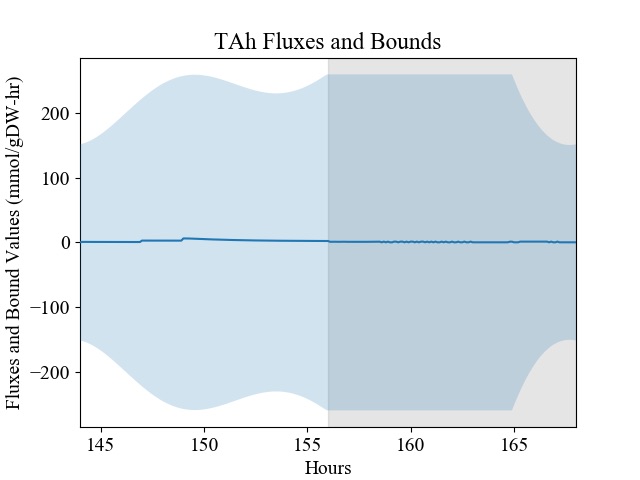

### Supplemental File 11

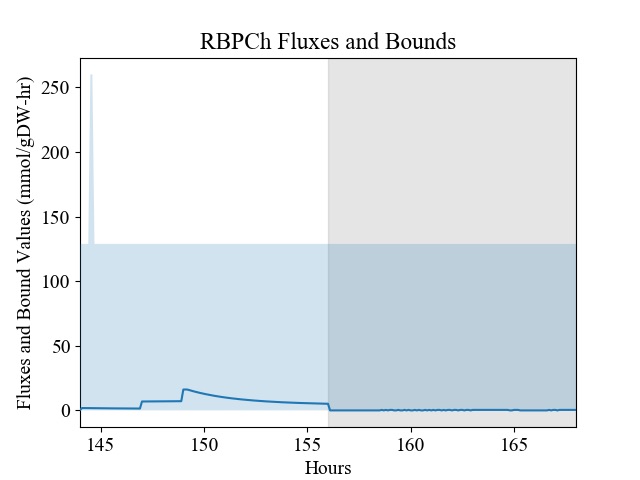

### Supplemental File 12

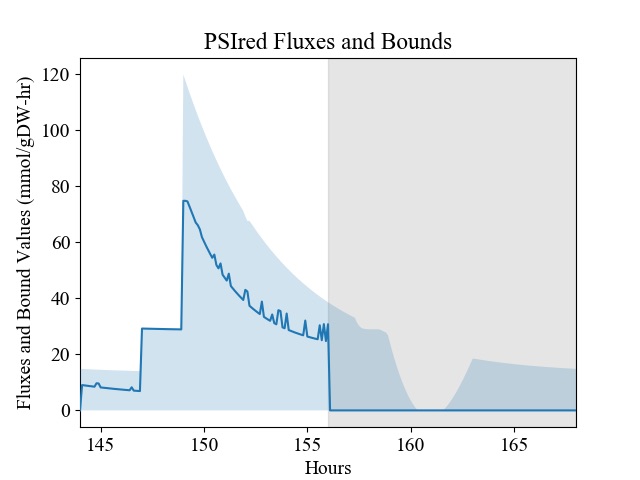

### Supplemental File 13

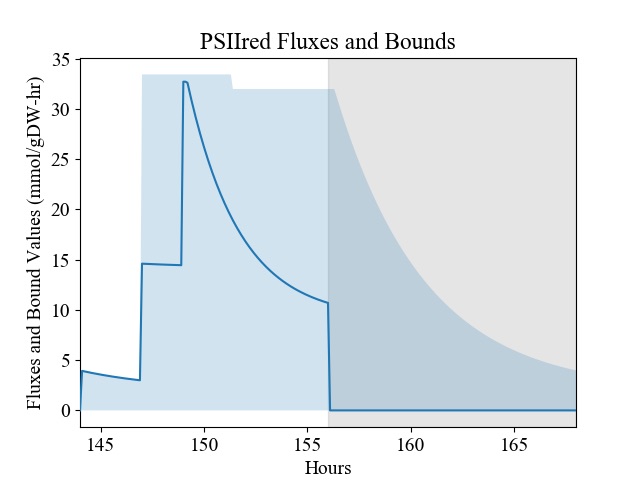

### Supplemental File 14

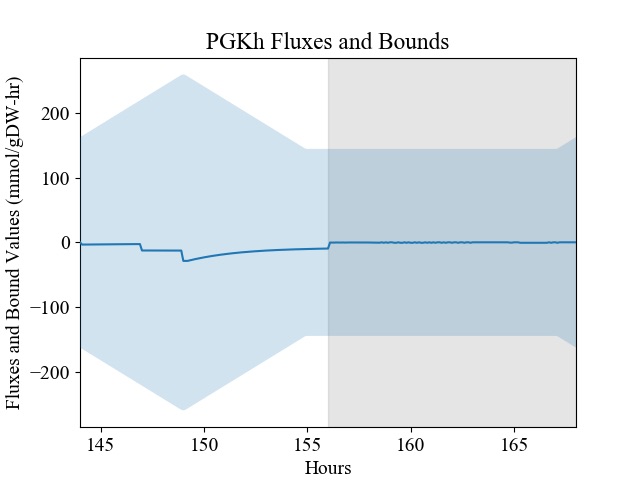

### Supplemental File 15

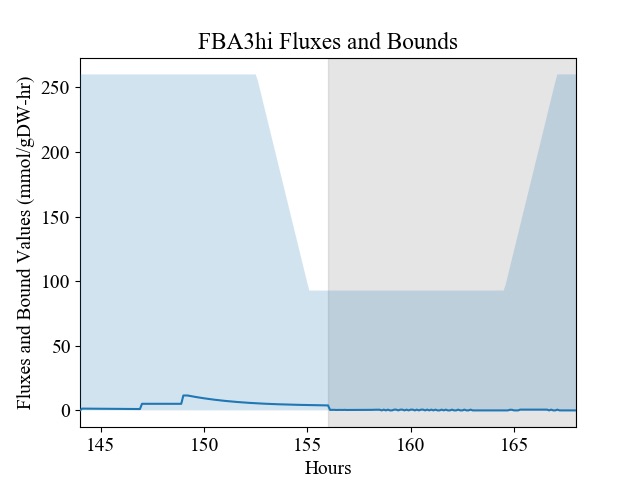

### Supplemental File 16

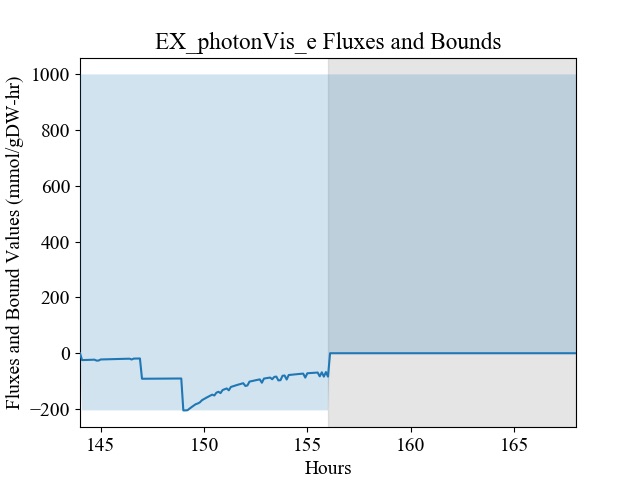

### Supplemental File 17

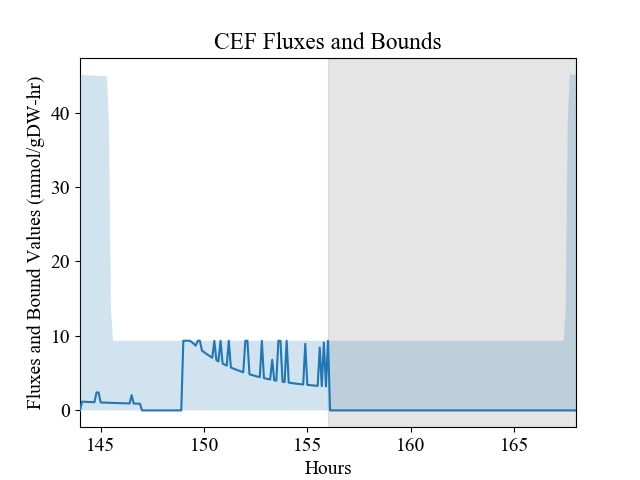

### Supplemental File 18

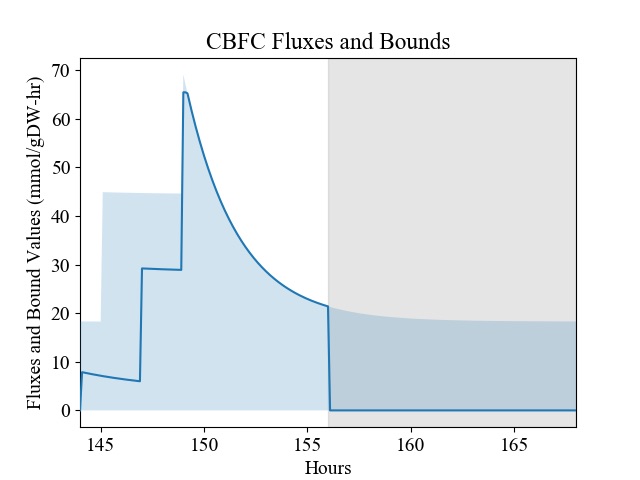
